# Viral Insulin/IGF-like Peptides Selectively Activate Host Insulin/IGF Signaling pathways during Grouper Iridovirus Infection

**DOI:** 10.1101/2024.06.13.598936

**Authors:** Aurelien Chuard, Kalaimagal Nesarajah, Khadija Danazumi, Kaitlin Reiners, Fa Zhang, Lev Levintov, Harish Vashisth, Richard Dimarchi, Emrah Altindis

## Abstract

Until our recent discovery of viral insulin/IGF-1-like peptides (VILPs), examples of viral mimicry were mostly limited to immunomodulatory proteins and growth factors. We previously showed that six viruses in the Iridoviridae family encode genes mimicking insulin and IGF-1. While VILP-carrying viruses are isolated from fish, the role of VILPs in host-pathogen interactions remain elusive. In this study, we used the Grouper Iridovirus (GIV), one of the VILP-carrying viruses, to examine the impact of the GIV-VILPs on the host IGF-1R/IR system during infection. Our structural analysis revealed that GIV-VILP is 35% identical to both zebrafish insulin and 35 IGF-1. We also showed that GIV-VILPs are early viral transcripts and resulting peptides are secreted during viral infection. Both single-chain (sc, resembling IGF-1) and double chain (dc, resembling insulin) forms of chemically synthesized GIV-VILPs can stimulate insulin receptor (IR) and IGF-1 receptor (IGF1R) phosphorylation. They could also stimulatepost-receptor signaling in both grouper kidney (GK) and zebrafish AB.9 cells as potently as insulin. Notably, supernatants obtained from GIV-infected cells could stimulate insulin/IGF system in a dose dependent manner. Interestingly, GIV-VILP was able to selectively activate the Akt/PI3K pathway while had no or minimal effect on Erk/MAPK pathway. Using either IR, IGF1R or dual inhibitors, we illustrated that inhibition of IR signaling suppressed GIV viral replication while inhibition of IGF1R enhanced viral replication. Consistent with selective signaling action of PI3K/Akt pathway, inhibition of the Akt reduced GIV viral replication however Erk pathway inhibition did not affect it. In summary, GIV-VILP is produced during GIV infection and manipulates host insulin/IGF signaling in a selective manner. Our findings reveal a previously unknown viral strategy in which viruses mimic a host hormone to exploit the host’s endocrine system. This discovery unveils a novel viral pathogenesis mechanism, broadens our understanding of viral mimicry, and opens a new avenue to better understand viral strategies to manipulate the host.

## INTRODUCTION

Viruses have evolved a variety of viral mimicry strategies to imitate host proteins, enabling them to manipulate host metabolism, cell cycle, cellular signaling cascades and evade the host immune response^1–3^. We recently showed that six viruses in Iridoviridae family have genes encoding viral insulin/insulin-like growth factor 1-like peptides (VILPs)^4^. In our previous studies, we chemically synthetized five of these VILPs and showed that all VILPs can bind to human insulin receptor (IR) and IGF-1 receptor (IGF1R) with varying potencies. While most of the VILPs, including all double chain (dc, resembling insulin) VILPs act as agonists of IR and IGF-1^4,5^, two VILPs act as antagonists of the IGF1R^6–8^. Specifically, single chain (sc, resembling IGF-1 containing a C-peptide) forms of Lymphocystis disease virus-1 (LCDV-1) and Mandarin fish ranavirus (MFRV) VILPs have very high affinity for the IGF1R. However, these two scVILPs inhibit IGF1R signaling. These unique characteristics are most likely acquired as a result of the distinct evolution of the VILPs in the viruses to facilitate viral pathogenesis, unlike the evolution of the host insulin/IGFs^9,10^.

Insulin/IGF system plays a key role in orchestrating cellular machinery. Insulin primarily regulates cell metabolism, including the glucose, protein, and lipid metabolism ^11,12^. On the other hand, the IGF-1 regulates cell proliferation, survival, and differentiation^13^. Because insulin and IGF-1 play key roles in human disease including diabetes and cancer, there has been extensive research over the last hundred years characterizing this system. However, our knowledge on the role of insulin/IGF system is still relatively limited for most organisms, including fish. While the main functions of fish insulin^14,15^ (regulating metabolism, development and feeding) and fish IGFs^16,17^ (proliferation, survival, differentiation and growth) are similar to that in mammals, fish show some important differences. For example, compared to mammals, fish express less IR and more IGF1R^18^ in their tissue^19–21^. Notably, our initial studies show that both forms of VILPs (sc and dc) have higher affinity for IGF1R compared to IR. Consequently, they are more potent on IGF1R^4,5,7^. In fish, IGF-1 is also involved in osmoregulation, as higher doses of IGF-1 stabilize plasma osmolality and sodium levels^22^. Furthermore, from a host-pathogen interaction perspective, both insulin and specifically IGF-1 play key roles in regulating both adaptive and innate immune responses in fish^23^.

In this study, we used Grouper Iridovirus (GIV) and grouper kidney (GK) cells to determine the role of VILPs in host-pathogen interactions. GIV is one of the six VILP carrying Iridoviridae family viruses with a large genome, ∼ 140K bp in length and contains 120 open reading frames^24^. It was recently isolated from spleen tissue of a diseased yellow grouper (*Epinephelus awoara*)^25^. GIV a serious pathogen in mariculture causing high mortality rates in finfish. Specifically, infection of cultured grouper (*Epinephelus* spp.) with GIV leads to heavy economic losses in fish aquaculture^26,27^. GIV infection induces cytopathic effects including cytoplasmic vacuolation, fusion, formation of cell aggregations and cell rounding in grouper cell lines^25,28^.

Our previous studies characterized chemically synthesized VILPs using mammalian systems. However, the role of VILPs on fish cells and its function in host-pathogen interactions remained unknown. Here, we address this gap by focusing on the effects of GIV infection and GIV-VILP on the fish insulin/IGF system during the infection. GIV-VILP share ∼30–50 % sequence homology with both human and zebrafish, insulins and IGF-1s (**Fig. 1A**). Using both chemically synthesized VILPs and GIV infection models, we showed that GIV-VILPs are actively secreted during the infection, stimulate IR/IGF1R and post-receptor signaling. Our findings demonstrate that insulin/IGF system is crucial for GIV replication. Manipulating insulin/IGF signaling during infection represents a novel viral pathogenesis mechanism. This previously unrecognized strategy, where viruses mimic host hormones to hijack cellular processes, unveils a new approach to understanding virulence mechanisms and may lead to new therapeutic targets for viral infections.

**Figure 1:**
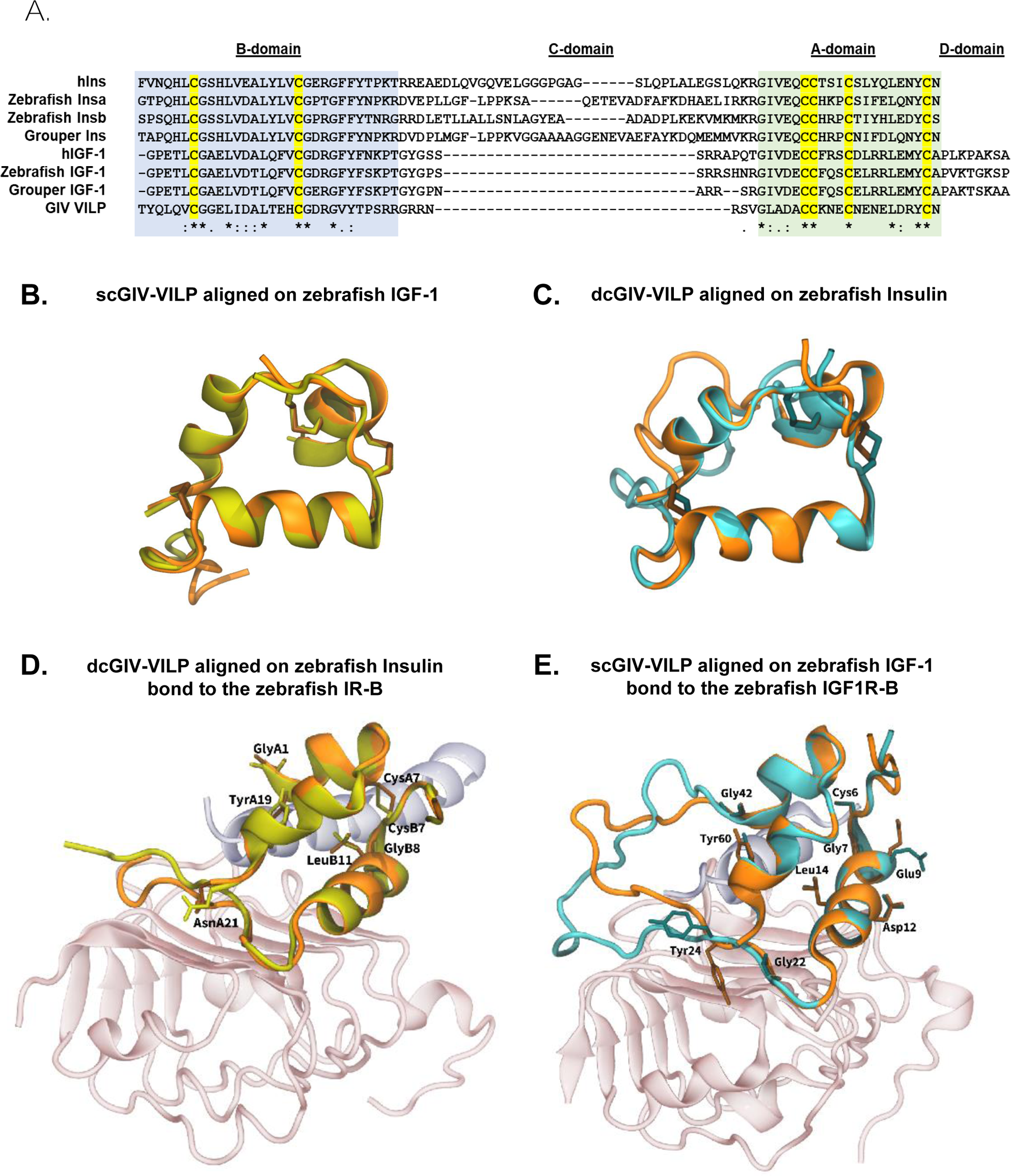
Grouper and zebrafish insulins and IGF-1s share highly similar amino acid sequence and maintain a conserved structure. and GIV-VILP. A-Sequence alignment of the B-, C-, and A-chains of insulin, IGF-1 from human, zebrafish and grouper compared to GIV-VILPs. Cysteine residues are highlighted in yellow. Asterisks signify identical residues, while periods and colons represent low and high degrees of similarity, respectively. B- Side-view snapshots of the dcGIV-VILP (orange cartoon) aligned over Zebrafish insulin (yellow cartoon) with conserved disulfide bridges highlighted as sticks. C- Side-view snapshots of the scGIV-VILP (orange cartoon) aligned over Zebrafish IGF-1 (cyan cartoon) with conserved disulfide bridges highlighted as sticks. Side-view snapshots of the primary binding site of (D) the zebrafish IR-B with DcGIV-VILP(orange cartoon) aligned over zebrafish insulin (yellow cartoon) and (E) of the zebrafish IGF1R-B with scGIV-VILP (orange cartoon) aligned over zebrafish IGF-1 (cyan cartoon) with conserved amino acid side chains highlighted as uniquely-colored sticks (main chain is shown for glycine). L1 and α-CT peptide are shown in pink and light-blue, respectively.

## RESULTS

### GIV-VILP exhibits significant homology to fish insulins and IGFs with a predicted insulin/IGF-like structure and distinct IGF1R binding

We previously showed that VILPs are 30-50% identical to human insulin and IGF-1^4,7^. Alignment analysis of GIV-VILP compared to human, zebrafish, grouper insulins and IGF-1s revealed that all six critical cysteine residues, essential for forming the intra- and interchain disulfide bonds required for insulin/IGF structures, are conserved among them (**Fig. 1A**). Unlike insulins’ long C-peptide, GIV-VILP possess a short seven amino acid C-peptide similar to IGFs. GIV-VILP exhibits 30% to 38% identity with the compared ligands (**Table 1**). Furthermore, the GIV-VILP sequence contains a 184 amino acid (aa) long extension domain composed of 18 repeated motifs (VAPST) which is potentially cleaved during processing of the ligand.

**Table 1:** Conserved residues among GIV-VILP and human, zebrafish, grouper insulins and IGF-1s.

We first performed structural modeling of both single and double-chain forms of the VILPs to explore the structural similarities among tertiary structures of VILPs with zebrafish insulin and IGF-1. We observed that VILPs exhibited insulin/IGF like structures, with the presence of two α-helices in the A-chain, an α-helix in the B-chain, and six conserved cysteines which formed three disulfide bridges (**Fig. 1B-C**). While dcGIV-VILP is predicted to have a similar configuration to zebrafish insulin (**Fig. 1B**), scGIV-VILP had a very similar structure to the zebrafish IGF-1 (**Fig. 1C**). The only difference in the scVILP was observed in the position of the C-peptide, which might be related to some distinct functions of the scGIV-VILP. the native Zebrafish IR/IGF1R binders, (**Fig. 1B-C**). We also completed structural analysis of the GIV-VILP binding to zebrafish IR (**Fig. 1D**) and IGF1R (**Fig. 1E**) compared to the native ligands labeling residues important for receptor binding. Our analysis revealed that dcGIV-VILP and zebrafish insulin have similar binding properties to the zebrafish IR and zebrafish IGF1R. Residues that are critical to IR binding such as GlyA1, TyrA19 and LeuB11 (**Fig. 1D**) and to IGF1R binding such as Leu14, Tyr24, and Tyr60 (**Fig. 1E**) are conserved and adopted similar positions upon receptor binding. On the other hand, C-peptide in the IGF1R showed significant differences in term of their positions. We previously showed that LCDV-1 scVILP which is a natural antagonist of the IGF1R and determined its binding to the IGF1R using cryoEM. We further compared the binding modes of dcGIV-VILP and scGIV-VILP to LCDV1-VILP. Both forms of the GIV-VILPs displayed similar configurations of the A- and B-chains to the corresponding chains in scLCDV1 with conserved amino acids adopting the same positions across these peptides (**Fig. S1**). Overall dcGIV-VILP displayed a binding mode that was more similar to IGF-1 than to scLCDV1 (**Figs. S1A, S2**). However, scGIV-VILP C-chain adopted a distinct configuration in comparison to the C-chain in IGF-1 (**Fig. 1C**) and scLCDV1 (**Fig. S1B)** indicating potential differences in function.

### GIV-VILPs is secreted during infection and it is an early gene product

To determine the cellular localization and potential secretion of GIV-VILP, we used a transfection model combined with imaging. To this end, we transfected GK cells with a pcDNA3.1+ plasmid expressing GIV-VILP fused to eGFP on its C-terminus. The transfected cells were fixed after 24 hours, mounted, and analyzed by confocal microscopy. We used a Golgi apparatus-specific antibody, anti-58K Golgi Protein, to visualize whether GIV-VILP-eGFP could be found in the secretory pathway. These imaging experiments showed that localization signals of GIV-VILPs overlapped with Golgi-associated signals. Correlation between the GFP signal and the 58K Golgi protein has been overall quantified with a Person correlation factor of 0.76, indicating that GIV-VILPs are in the secretory pathway (**Fig. 2A**). To further examine whether VILP is a secreted protein, we used Brefeldin A, a potent inhibitor of the Golgi apparatus. We transfected GK cells with the same construct and used Brefeldin A inhibitor for 6 hours after the 24-hour transfection. Western blot analysis demonstrated a significant increase in intracellular GIV-VILP-eGFP signal when comparing samples with or without Brefeldin A, confirming the processing of GIV-VILP for secretion (**Fig. 2B**). SignalP 5.0 predicted that GIV-VILP (*UniProt Q5GAI4*) possesses a signal peptide before B chain that will allow the peptide to be targeted towards the secretory pathway (**Fig. S3**).

**Figure 2:**
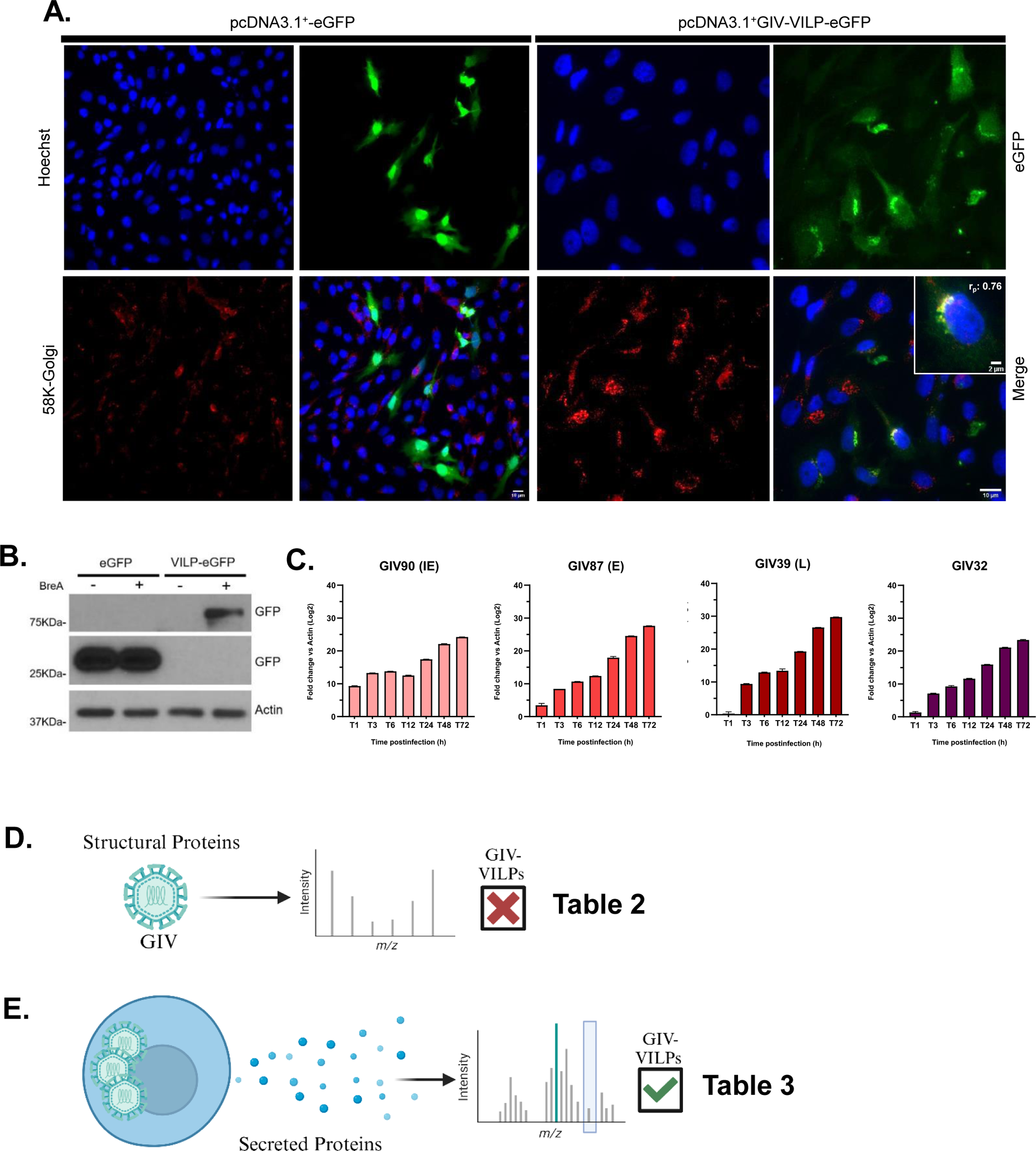
GIV-VILPs are nonstructural peptides secreted through the Golgi apparatus during GIV infection. A- Immunofluorescence of GK cells transfected with either pcDNA3.1+-eGFP plasmid or pcDNA3.1+-GIV-VILP-eGFP. pcDNA3.1+-GIV-VILP-eGFP expressing cells with GIV-VILP-eGFP in green colocalize with Golgi apparatus highlighted by 58K-Golgi antibodies in red, nucleus was stained with Hoechst (blue). B- Western blot analysis of GIV-VILP-eGFP proteins levels after 6h treatment with Brefeldin A (10µg/mL). (C) Viral transcripts GIV90, GIV87, GIV39, and GIV32 (GIV-VILP) were quantified by RT-qPCR after RNA extraction and cDNA synthesis from GK infected cells (MOI 1) at different hours post infection (hpi). Data are expressed as means ± SDs (n = 3). B-Actin is used for normalization. MOI:1. (D) Scheme of the mass spectrometry analysis of virions’ structural proteins, refer to Table 1. (E) Scheme of the mass spectrometry analysis of secreted proteins from the infection of GK cells by GIV virus, refer to Table 2.

**Table 2:** Structural proteins of GIV virions. Proteins identified from purified GIV virions by LC/MS mass spectrometry. Predicted functions, protein sizes, accession numbers, and associated orthologs are listed.

Active transcription of GIV-VILP and its kinetics were subsequently examined during viral replication. Infected GK cells at a Multiplicity of infection of 1 (MOI 1) were collected at 1, 3, 6, 12, 24, 48, and 72 hours post-infection (hpi). RNA extraction and RT-qPCR were performed to quantify viral transcripts. Based on prior kinetics experiments conducted on the Singapore Grouper Iridovirus (SGIV), a sister virus to GIV, we selected GIV90, GIV87 and GIV39 (major capsid protein) as immediate early, early, and late gene controls, respectively. Our results validated this selection and all control genes exhibited an exponential increase in the transcription mirroring viral replication (**Fig. 2C**). GIV32, encoding GIV-VILP, was transcribed as early as 3 hours post-infection, similar to GIV87 transcripts, allowing us to define GIV32 encoding GIV-VILP as an early viral gene (**Fig. 2C**).

### GIV-VILP is a secreted protein during infection rather than a structural protein

We purified GIV virions and analyzed the viral particle protein composition using LC/MS mass spectrometry (**Fig. 2D and Table 2**). We identified 20 structural proteins, including major capsid protein (GIV39), in all three independent experiments. Using BlastP, we determined the orthologs of these proteins and predicted functions (**Table 2**). Notably, GIV01 (Rho_N domain-containing protein), GIV36 (Surface protein), GIV17 (Protein kinase domain-containing protein), GIV 76 (Methyl-accepting transducer domain-containing protein) and GIV44 (Putative tyrosine protein kinase) and GIV39 (Major capsid protein) were the top hits identified in all three experiments with the highest number of peptides identified in the spectrum. Intriguingly, GIV contained two structural protein kinases, an RNAse and a collagen like protein. These results also indicate that GIV-VILP is not a structural protein of GIV virions.

Following the structural analysis, we analyzed the secretome of the cells infected with the GIV (**Fig. 2E**). Samples were collected 24 hours post-infection, and secreted proteins were precipitated and analyzed using LC/MS mass spectrometry. In total, we identified 10 secreted GIV proteins in three independent experiments. Notably, GIV-VILP (GIV32) was one of the most abundant proteins. In addition, ubiquitin-like domain-containing protein (GIV57), sema domain-containing protein (GIV87) and collagen-like protein (GIV63) were other most abundant proteins (**Table 3**). Notably, there were two others viral proteins mimicking another host hormone, fibroblast growth factors (FGF), FGF-10 (GIV 79) and FGF-22 (GIV 78). Finally, we identified a second collagen-like protein (GIV62) in the secretome, in addition to the one identified in the structural proteins (GIV21). Ig-domain containing protein (GIV 14) was another interesting protein, potentially involving in immune evasion. Overall, these results demonstrate active secretion of the VILPs during the GIV viral cycle.

**Table 3:** Protein composition of GIV secretome. TCA-precipitated proteins were identified by LC/MS mass spectrometry. Predicted functions, protein sizes, accession numbers, and associated orthologs are listed.

### Chemically synthesized GIV-VILPs are active ligands of grouper IGF1R and/or IR

Previous studies characterized VILPs using murine and humanized cell models^4,7–9^: however, the impact of VILPs on their native hosts, fish cells and fish IR / IGF1R signaling has not been studied. To address this gap, we first chemically synthesized both single-chain (sc, IGF-1 like) and double-chain (dc, insulin like) forms of GIV-VILP as previously described^4,7^ Using GK cells, we tested whether chemically synthesized GIV-VILPs can stimulate host IGF1R/IR autophosphorylation and activate post-receptor signaling. We conducted dose-response experiments (1 nM, 10nM, 100nM for 15 minutes) with Akt phosphorylation indicating PI3K/Akt pathway activation and of Erk phosphorylation indicating Ras/MAPK pathway activation.

Fish cells are known to be more susceptible to IGF-1 rather than insulin^16–19^. Consistent with the literature, GK cells were very sensitive to IGF-1, and receptor phosphorylation was strongly induced by IGF-1 compared to insulin (**Fig. 3A**). On the other hand, scGIV-VILP and DcGIV-VILPwere comparable to insulin in stimulating receptor phosphorylation (**Fig. 3A**). The PI3K pathway was stimulated by all the ligands and followed the order of stimulation observed with the receptor signaling, with stronger stimulation by IGF-1 compared to the other three ligands (**Fig. 3A**). However, we did not observe a similar dose response in the phosphorylation of Erk (MAPK pathway).

**Figure 3:**
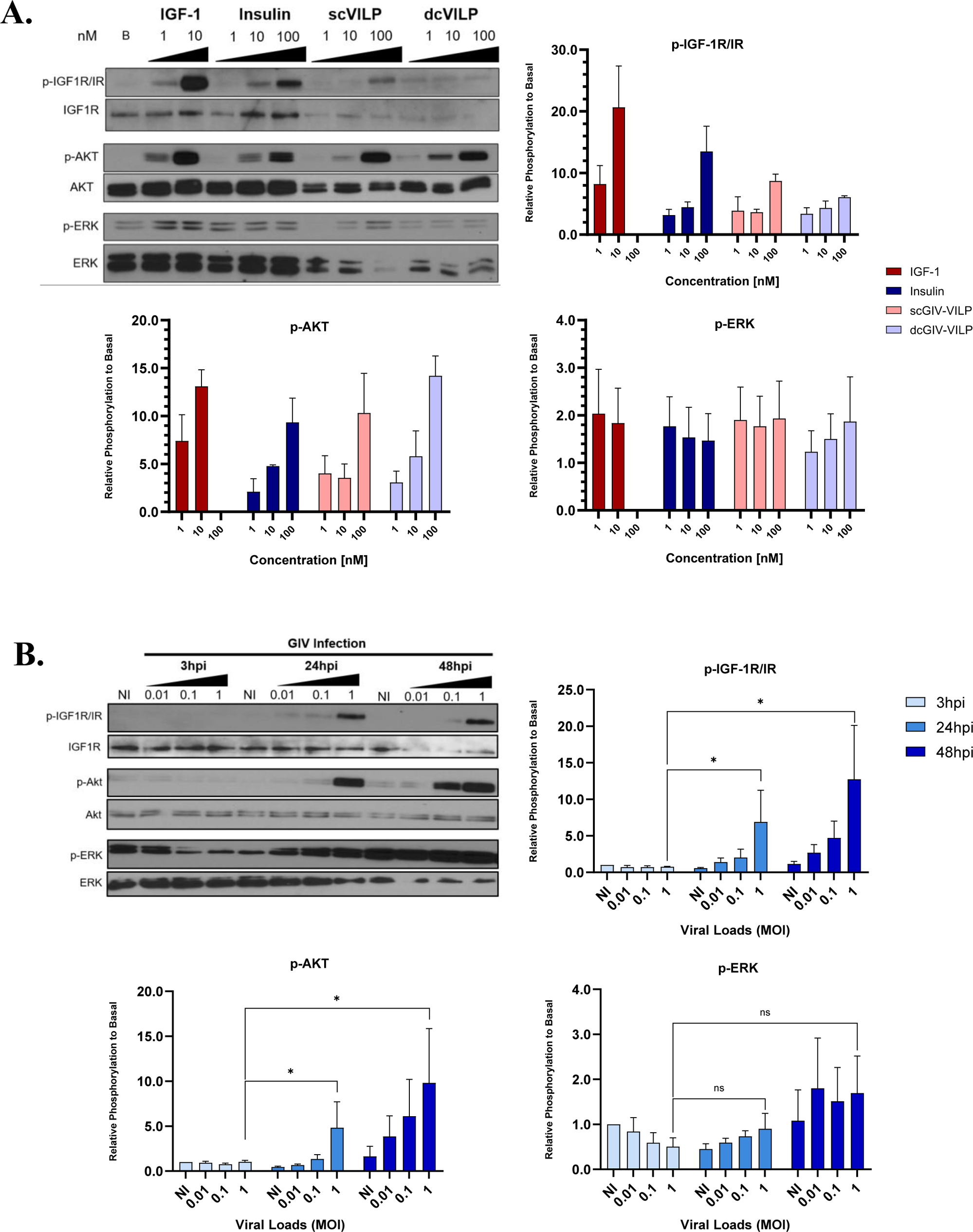
GIV-VILPs are active ligands of IGF1R/IR in GK cells, resulting in the selective activation of Akt downstream signaling. Phosphorylation of IGF1R/IR, Akt and Erk of GK cells was monitored by Western blot **(A)** after stimulation with IGF-1, insulin, scGIV-VILP or dcGIV-VILP at different concentrations (nM). and (B) GIV infection at different viral loads (MOI 0.01/ 0.1/ 1) and at different time points (3hpi, 24hpi and 48hpi). All experimental conditions are normalized to basal conditions assuming1 as the constant value for basal. Exposure times ranged from 10 seconds to 1 minute. All experiments were performed in triplicate and statistical analysis were performed using unpaired t-test (*p < 0.05; **p < 0.01)

**Table 4:** List of oligonucleotides used in this study.

### GIV Infection Induces Receptor Phosphorylation and Selective pAkt Signaling

To assess the VILP function during infection, we infected serum-starved GK cells with varying doses of GIV (MOI 0.01, 0.1, 1) at three different time points: 3 hpi, 24 hpi, and 48 hpi. Upon completion of the experiment, the supernatant from infected cells was transferred to serum starved GK cells for signal stimulation analysis. At 3 hpi, GIV infection did not stimulate insulin/IGF signaling (**Fig. 3B**). This is consistent with the GIV-VILP gene expression profile, confirming that it is not an immediate early gene. However, at 24 hpi and 48 hpi, in a viral load-dependent manner, we observed strong receptor phosphorylation as well as a biased post-receptor signaling phosphorylation of Akt. Interestingly, phosphorylation of Erk was not correlated with viral replication (**Fig. 3B**) and this is consistent with our earlier observation when we used chemically synthesized VILPs (**Fig. 3A**) These results indicate that GIV-VILP produced during infection can bias host insulin/IGF signaling specifically towards the PI3K/Akt pathway rather than MAPK/Erk pathway.

### Zebrafish cells are susceptible to GIV infection, and GIV-VILPs stimulate zebrafish insulin/IGF signaling

Because most of the immune system related components, cells and signaling pathways are conserved between fish and mammals, zebrafish serves as a model to study host-virus interactions^29^. Several previous studies have used zebrafish model to study pathogenesis for both fish and human viruses^30–33^. Notably, the zebrafish was used to characterize two other Iridoviridae species^34,35^. However, the susceptibility of zebrafish to GIV and the effects of GIV-VILPs remain unknown. To determine whether zebrafish can be used as a model system to study GIV infection and to assess the reproducibility of our results in GK cells in a different fish model, we utilized a zebrafish fibroblast cell line (AB.9). We first showed that cytopathic effects are induced by GIV after 3 days, with cells undergoing lysis after 7 days (**Fig. S4**). To examine viral kinetics, and the expression of GIV-VILP, we assessed the same viral genes (GIV90, GIV87, GIV39) as controls as described above. Consistent with our findings in grouper cells, GIV-VILP functions as an early gene in zebrafish model as well (**Fig. 4A**). We subsequently examined the effects of GIV-VILPs on signaling. Among the ligands tested, IGF-1 exhibited the highest potency, while both forms of the VILPs showed comparable abilities to stimulate receptor phosphorylation as insulin. Although we were unable to assess Erk phosphorylation due to elevated basal levels, all ligands induced Akt phosphorylation (**Fig. 4B**). Overall, these results indicate that zebrafish cells are susceptible to GIV infection and it could be used as a model system. Furthermore, GIV-VILP is an important early gene that is expressed both in grouper and zebrafish cells to manipulate host insulin/IGF signaling.

**Figure 4:**
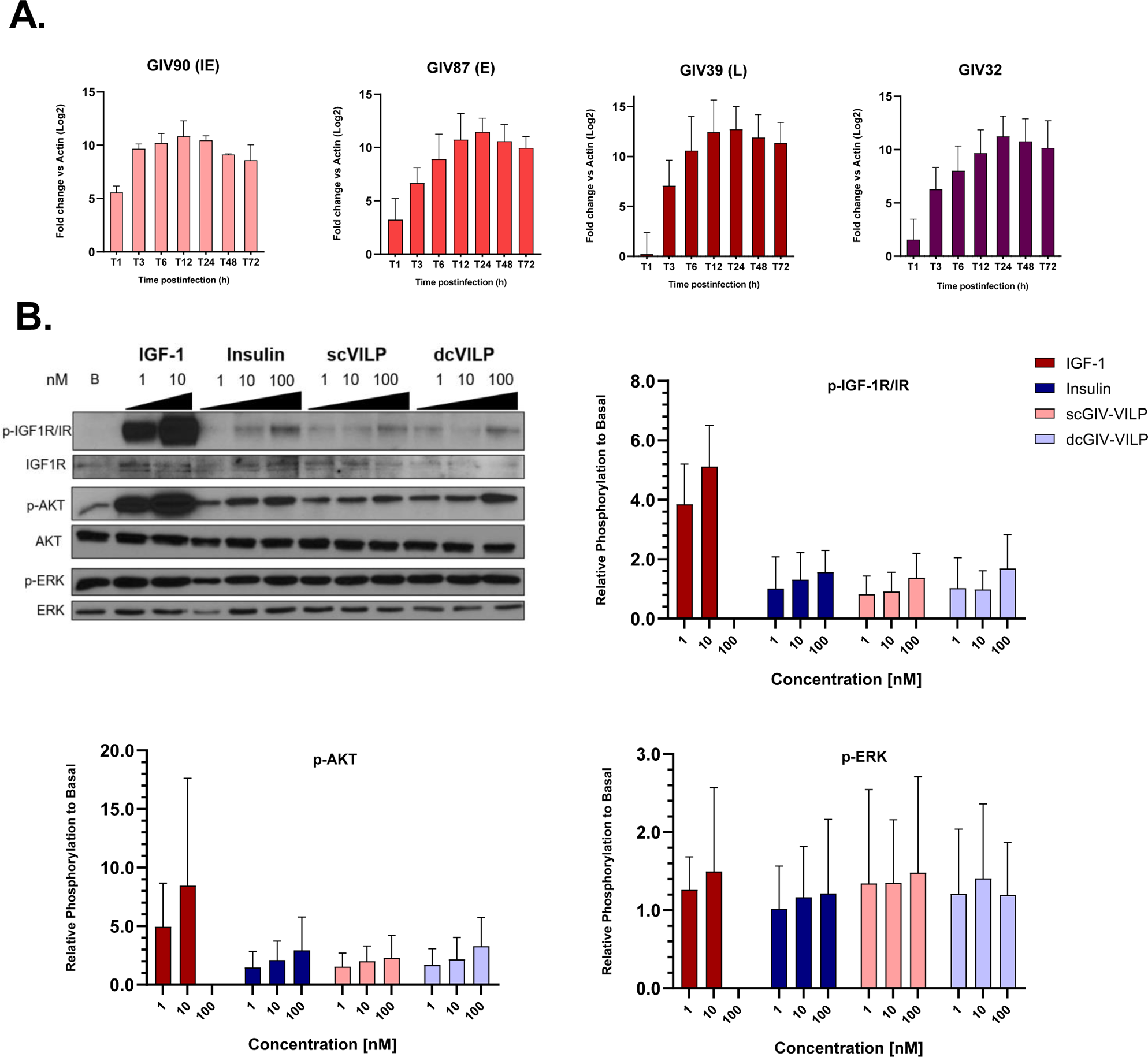
GIV-VILPs are active ligands of IR /IGF1R in AB.9 zebrafish cells, resulting in the selective activation of Akt downstream signaling. A. Phosphorylation of IGF1R/IR, Akt, and Erk in zebrafish AB.9 cells were monitored by Western blot after stimulation with IGF-1, insulin, scGIV-VILP, or dcGIV-VILP at different concentrations (nM). All experimental conditions are normalized to basal conditions, with 1 used as the value for basal. Exposure times ranged from 10 seconds to 1 minute.

### Inhibiting IR or Akt Activity Decreases GIV Replication, while IGF1R Inhibition Promotes It and Erk Inhibition Has No Effect

The presence of a VILP in the GIV genome indicates the involvement of the insulin/IGF syste in viral replication. To determine the roles of IR, IGF1R and two related post-receptor signaling pathways (PI3K/Akt and MAPK/Erk) in GIV replication, we used specific inhibitors and competitive ligands of these receptors. We first used Picropodophyllin (PPP), a tyrosine kinase inhibitor specific to the IGF1R during infection Surprisingly, inhibition of IGF1R by PPP (2μM) resulted in a substantial increase in viral replication of 140% (**Fig. 5A**). Subsequently, we tested the impact of an IR-specific peptide antagonist S961 and showed that S961 decreased GIV replication by 54% at the lowest dose tested (10 nM) (**Fig. 5B**). Finally, BMS-754807, an inhibitor that targets both IR and IGF1R increased replication by 118% at concentration 0.1nM (**Fig. 5C**). Although, magnitude of these increases was lower than IGF1R inhibitor’s effect, the relatively higher expression of IGF1R in these cells suggest that BMS-754807 may exert an effect similar to IGF1R inhibitor. These results demonstrate that while IR signaling promotes viral replication, the same does not hold true for IGF1R signaling. In fact, activation of the IGF1R pathway appears to be detrimental to viral replication.

**Figure 5:**
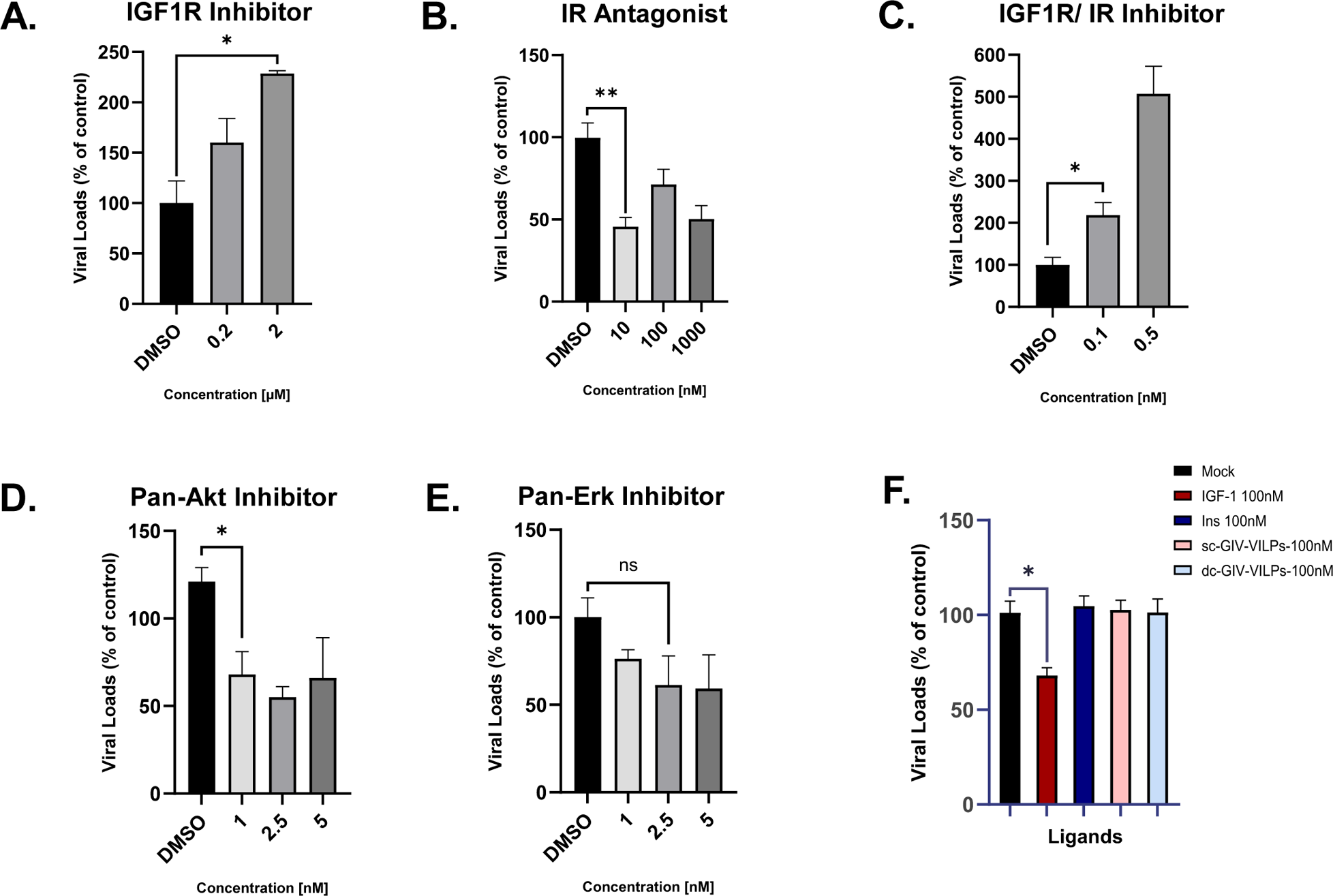
Inhibiting IR or Akt Activity Decreases GIV Replication, While IGF1R Inhibition Promotes It and Erk Inhibition Has No Effect. Quantification of GIV replication rates obtained after 12 or 24 hours of infection. (A) Quantification of GIV replication after 12 hours stimulation with either Insulin, IGF-1, scGIV-VILP or dcGIV-VILP. (C) Quantification of GIV replication after 24h treatment by IGF-1-R or IR inhibitors. (D) Quantification of GIV replication after 24h treatment by Pan-Akt or Pan-Erk inhibitors. All quantifications were performed in triplicate and statistical analysis were performed using unpaired t-test (*p < 0.05; **p < 0.01)

Our signaling experiments demonstrated that GIV-VILPs were potent stimulators of Akt phosphorylation while had almost no effect on Erk phosphorylation (**Fig. 3A**), indicating the importance of biased PI3K pathway activation for GIV replication. Consistently, inhibition of the Akt pathway using MK-2206 led to a 32% decrease in viral replication (**Fig. 5D**). Whereas inhibition of Erk using MK-2206 had no significant effect GIV replication (**Fig. 5E**).

Finally, we tested the direct impact of IGF-1, insulin, scGIV-VILP, or DcGIV-VILPon GIV replication. GK cells were inoculated with GIV at a MOI of 1 in serum-free media for 1 hour. Subsequently, they underwent repeated stimulation every 3 hours with the specified ligands at a concentration of 100nM for a total duration of 12 hours. GIV replication was quantified at the end of the experiment. Consistent with our IGF1R inhibition experiments, IGF-1 stimulation led to a 34% decrease in GIV replication, whereas stimulation with insulin, scGIV-VILP, and dcGIV-VILP did not decrease the replication (**Fig. 5F**).

## DISCUSSION

Over millions of years, viruses have co-evolved with their hosts, resulting in intricate interactions and adaptations with their associated host. Viral mimicry mechanism, wherein viruses mimic host proteins, arise from host-virus interactions, aiding viruses in subverting or manipulating cellular processes to their advantage^36^. Most of these viral proteins’ mimic hosts’ growth factors. For example, poxviruses are known to encode epidermal growth factor (EGF) mimics^37^. Likewise, the first growth factor identified for vaccinia virus was vaccina growth factor^38^. Similarly, viruses are evolved to evade host immune response-associated proteins using mimics. For instance, Kaposi’s sarcoma-associated herpesvirus (KSHV), Cytomegalovirus (CMV) and Herpesvirus Saimiri gene 13 (HVS13) viruses carry cytokine mimicking IL-6, IL-10, and IL-17, respectively ^39–42^. In addition, similarities between viral proteins and human proteins are linked to autoimmunity through molecular mimicry For example, recent studies identified a potential link between Epstein–Barr virus (EBV) and the onset of multiple sclerosis (MS) via this mechanism. Researchers identified cross-reactive antibodies that bind to both EBV nuclear antigen EBNA1 and glial cell adhesion protein in the central nervous system^43^. Epidemiological evidence further supports this connection between EBV infection and MS autoimmunity^44^. Our findings reported in this study have broadened our knowledge of viral mimicry mechanisms. Here, we demonstrate that certain viruses developed insulin/IGF mimicry mechanisms to directly act on the IGF1R and IR, thereby modulating post-receptor signaling in a selective manner and modulate related cellular mechanisms.

In this study, we used both grouper and zebrafish cells and showed that GIV-VILP is transcribed as an early gene and translated during infection. VILPs are secreted proteins and are not identified in virions. GIV-VILPs have capability to selectively stimulate host IR/IGF1R signaling, favoring activation of PI3K pathway while not inducing activation of the MAPK/Erk pathways. Using inhibitors, we demonstrated that PI3K/Akt pathway is indeed crucial for viral replication because inhibition of the Akt pathway, rather than Erk, leads to a decrease in GIV replication. Notably, our findings also revealed that inhibiting the IGF1R enhances GIV replication, whereas inhibiting the IR decreases GIV replication. Consistent with this observation, direct stimulation by IGF-1 decreases GIV replication, whereas, insulin, scGIV-VILP, and dcGIV-VILP do not have any negative effects on replication.

The PI3K/Akt pathway is highly conserved among vertebrates and regulates host metabolism. Previous research has shown that PI3K pathway is an important target of viruses. For example, HSV-1 entry relies on the PI3K activity as its inhibition decreases HSV-1 entry through an unknown mechanism^45^. PI3K regulation of the nuclear translocation of Serine/Threonine (SR) proteins that lead to pre-mRNA splicing events impacts HIV-1 replication if this pathway is inhibited^46^. In addition, viruses target PI3K pathway to inhibit apoptosis. For example, HSV-1 and HCMV encode US3 and Major immediate-early proteins (MIEP), respectively, that constitutively mimic Akt activity and promoting cell survival by preventing apoptosis^47,48^. Lastly, inhibiting PI3K/Akt during human immunodeficiency virus (HIV-1) infection was sufficient to completely halt single-round HIV-1 infection^49^. As previously mentioned, these regulations modulate key regulatory or adaptative proteins to regulate the PI3K/Akt pathway or glucose metabolism. However, specific viral insulin/IGF-1-like peptides that directly interact with IR and/or IGF1R receptors to stimulate these PI3K/Akt pathways were not identified until this study.

Insulin and IGF-1 bind to and activate their respective receptors, the IR and IGF-1-R, stimulating a conformational change in the receptor structure that initiates receptor autophosphorylation^50,51^. Our research demonstrates that inhibiting IGF1R benefits GIV replication, while inhibiting IR hinders it. While these receptors are structurally and functionally similar to each other, they have some significant differences in terms of their down-stream effects. In this study, the inhibition of IGF1R stimulated GIV replication. Conversely, IR inhibition results in a decrease of GIV replication. This highlights the importance of the IR signaling and IR-related downstream pathways for the GIV replication. Indeed, IR associated phosphorylation patterns compared to those of IGF1R, are strongly linked to the regulation of mTORC1 signaling as well as PI3K signaling modulating cell metabolism, as well as autophagy and apoptosis^12,52^. This difference in signaling patterns between the two receptors could explain the difference observed in our study. GIV may selectively stimulate PI3K/Akt phosphorylation through IR to enhance nutrient uptake and activate cell metabolism providing the necessary cellular resources for viral replication. By avoiding the direct stimulation of IGF1R and MAPK/Erk pathway, the virus prevents cell from entering a proliferative state, allowing cellular machinery to remain available for optimal viral replication. Supporting this hypothesis, we previously showed that MFRV-scVILP and LCDV1-scVILP are the first specific peptide inhibitors of IGF1R signaling^7,53^. Our findings show that IGF1R inhibition increase GIV replication which may be a common virulence mechanism for some VILP carrying viruses. Interestingly another group also showed that LCDV1-scVILP functioned as an inhibitor of ferroptosis, resulting in the prevention of cell death, which could be leveraged by the LCDV virus to increase its replication^54^.

Our structural modeling indicates some potential directions to better understand, GIV-VILP’s unique selective function on PI3K pathway. For example, the differences between IGF-1, LCDV-1 and scGIV-VILP for IGF1R binding (**Fig. S1B**) could be attributed to differences in the length of the C-chain, since the C-chains in IGF-1 and LCDV1-scVILP are five and three amino acids longer than the C-chain in scGIV-VILP, respectively, thereby potentially providing IGF-1 and LCDV1-scVILP additional residue contacts for interacting with the important receptor binding sites, L1 domain and the αCT peptide. This difference in C-peptide and its unique interaction with the receptors may be related to the selective signaling.

In humans, the dysfunction of insulin/IGF system is associated with various diseases including diabetes, obesity, cancer, growth impairments, and neurodegenerative diseases^55–58^. Our findings on the VILPs unveils a new mechanism in which insulin/IGF-1 system is modulated by viruses. Iridoviridae virus family members are known to infect ectothermic vertebrates^59^. However, we and others previously identified the DNA of VILP-carrying viruses in the gut microbiome and human bloodstream^4,60^. Interestingly, GIV was recently identified as one of the most abundant viruses in the obese patients’ gut virome compared to the lean controls^60^.The potential impact of VILP-carrying viruses in human health has not yet been investigated. However, there are only 11,700 viral genomes available in the NCBI database (June 2024). Nonetheless, Anthony et al. estimated that there might be over 320,000 mammalian viruses waiting to be discovered^61^ (P. In 2014, we identified only three VILPs; by 2019, this number increased to six, thanks to new sequencing projects. Given that insulin/IGF signaling plays a central role in regulating almost all functions in mammals, we believe that insulin/IGF mimicry may not be specific to fish viruses. Furthermore, we posit that it is possible to identify VILPs in human and/or mammalian viruses.

To our knowledge, this is the first study to define a viral insulin/IGF mimic that directly interacts with the host’s receptor and selectively stimulates post-receptor signaling. This novel viral pathogenesis mechanism significantly advances our understanding of viral infections and their impact on host physiology. Furthermore, it opens a new avenue to define the concept of viral mimicry, encompassing these viral hormones. These findings may benefit therapeutic interventions targeting these pathways, particularly in the fish industry, by exploring the development of IR antagonists or Akt inhibitors in lower doses to mitigate viral replication. Overall, our findings underscore the importance of considering insulin/IGF mimicry as an important factor in the pathology of infectious diseases, promising deeper insights into virus-host interactions and their implications for health and disease.

---

**Figure S1:** Side-view snapshots of the primary binding site of the zebrafish IGF1R-B with (A) GIV-dcVILP(orange cartoon) aligned over scLCDV-1 (green cartoon) and (B) with scGIV-VILP (orange cartoon) aligned over scLCDV-1 with conserved amino acid side chains highlighted as uniquely-colored sticks (main chain is shown for glycine). L1 and α-CT peptide are shown in pink and light-blue, respectively.

**Figure S2:** Side-view snapshot of the primary binding site of zebrafish IGF1R-B with dcGIV-VILP (orange cartoon) aligned over zebrafish IGF-1 (cyan cartoon) with conserved amino acid side chains highlighted as uniquely-colored sticks (main chain is shown for glycine).

**Figure S3: SignalIP analysis of the GIV-VILP amino acid sequence.**

SignalIP 5.0 analysis of the GIV-VILP amino acid sequence reveals a likelihood of 0.9423 for the presence of a signal peptide. The signal peptide suggests a potential cleavage after the threonine at position 19.

**Figure S4: Microscopic view of GK and AB.9 cells infected with GIV.**

Grouper kidney cells **(A)** and AB.9 cells **(B)** infected and non-infected with GIV, were observed4- or 7-days post infection, respectively.

## METHODS

### Peptide Alignment and Peptide Structure Prediction

Identified Viral Insulins-like Peptides used in this study (GIV-VILP) were aligned with corresponding Insulin and IGF-1 sequences from different organisms of interest, such as human (*Homo sapiens*); zebrafish (*Danio rerio*) and grouper (*Epinephelus coioides*) using ClustalW interface.

### Structural modeling

We used MODELLERv10.5 to generate the tertiary structures of dcGIV and Zebrafish insulin based on the template of the human Insulin structure (PDB: 6PXV), and scGIV and Zebrafish IGF-1 based on the template of the human IGF-1 structure (PDB: 6PYH). Additionally, we modeled truncated Zebrafish IR (μIR) and truncated Zebrafish IGF1R (μIGF1R) using the human IR (PDB: 6PXV) and the human IGF1R (PDB: 6PYH), respectively, as templates. Specifically, μIR was composed of the L1 domain (residues 1-155) and the αCT peptide (residues 691 to 717), and the μIGF1R was composed of the L1 domain (residues 1-155) and the αCT peptide (residues 691 to 705). We first modeled any missing residues (e.g., residues 38 through 40 in IGF-1) in the templates using the MODELLER software before constructing the tertiary structures of VILPs, Insulin, and IGF-1. We preserved the disulfide bonds across all peptide structures and maintained the double-chain configuration of peptides wherever necessary during model building. We generated 200 models of each peptide and the L1 domain, and selected the best model based on the lowest discrete optimized protein energy (DOPE) score. We superimposed the L1 domain and the αCT peptide from the Zebrafish IR and IGF1R on the human truncated IR (PDB: 6PXV) and IGF1R (PDB: 6PYH), respectively, and further superimposed peptide ligands (VILPs and Zebrafish Insulin/IGF-1) on the human Insulin/IGF-1 initially present in the human IR/IGF1R structures to obtain a complexed structure of each ligand using the PyMOL and VMD software tools.

### RNA Extraction and RT-qPCR

RNA from GK or AB.9 cells was purified using the Direct-zol RNA Miniprep Kit (Zymo #R2050) according to the manufacturer’s instructions. Reverse transcription was performed using 2µg of RNA with the Maxima First Strand cDNA Synthesis Kit (ThermoFisher #K1641), following the manufacturer’s instructions. RT-qPCR was performed using the SYBR™ Green PCR Master Mix (ThermoFisher #4309155) and specific oligonucleotides for each targeted transcript (**Table 3**). Samples were amplified on the QuantStudio 3 Real-Time PCR System (ThermoFisher #A28567). Quality control and visualization were performed using the associated QuantStudio™ Design & Analysis Software. RT-qPCR quantifications were conducted using the 2−ΔΔCT method, employing the relative Ct abundance of cellular and viral transcripts.

### Plasmids

All plasmid constructions were derived from pcDNA3.1+ (REF). Gene cloning was accomplished through enzymatic restriction, followed by ligation and transformation processes using One Shot® TOP10 Electrocomp™ E. coli (Invitrogen #C4040-50). Complete cloned plasmid sequences and accurate alignments were analyzed by Sanger sequencing performed by the Azenta Life Science platform. The sequences of oligonucleotides used for plasmid construction are listed in **Table 3**.

### Cells and viruses

Grouper Kidney (GK) cells were maintained in Leibovitz-15 medium (L-15) containing 10% Fetal Bovine Serum (Thermoscientific #A5256801), 1% L-Glutamine (Sigma-Aldrich #G3126), and 1% Penicillin-Streptomycin (Sigma-Aldrich #P4333) at 28°C without CO2.

Zebrafish caudal fin cells (AB.9) were purchased from ATTC (CRL-2298 ™). They were maintained in DMEM, high glucose (Thermofisher #11965092), 15% Fetal Bovine Serum, 1% L-Glutamine, and 1% Penicillin-Streptomycin. Grouper Iridovirus (GIV) was amplified on GK cells, and titration was performed by plaque assay on GK cells.

GIV viral stocks were prepared by infecting GK cells until the cellular monolayer was disrupted. Cells and viruses were collected by centrifugation at 12,000 x g for 30 min at 4°C. The cell pellet containing the virus was then resuspended in culture medium and ultrasonicated on ice (5 rounds of 1-minute intervals). Cell debris was removed by centrifugation at 4,000 x g for 20 min at 4°C. The supernatant was layered onto a 35% sucrose cushion and centrifuged at 210,000 x g for 1 hour at 4°C. The resulting pellet, resuspended in TN buffer (50 mM Tris-HCl [pH 7.4], 150 mM NaCl), was overlaid on 30%, 40%, 50%, and 60% (w/v) sucrose gradients and centrifuged at 210,000 x g for 1 hour at 4°C. The lowest band (50% sucrose) was individually aspirated and spun down at 100,000 x g in PBS solution. Aliquots of pure GIV virions are then stored at −80°C.

### Transfection and Immunostaining

Cells were seeded in P12 plates with coverslips previously placed at the bottom of each well. Transfections were performed using Lipofectamine™ 3000 Transfection Reagent (ThermoFisher #L3000001) following the manufacturer’s instructions. After 24 hours post-transfection, cells were fixed with 4% Paraformaldehyde and stored at 4°C until further processing. Immunostaining of the fixed samples was performed at room temperature. Firstly, samples were permeabilized and blocked in PBS-BSA (0.5%)-Triton (0.1%) for 30 minutes. After the blocking step, 58K Golgi Protein Antibody (Novus Biological #NB600-412SS) was diluted to 1/200 in PBS-BSA (0.5%)-Triton (0.1%) before applying to samples for 1 hour. Samples were washed 3 times with PBS-Triton (0.1%). The secondary antibody Alexa Fluor® 594 anti-rabbit IgG was diluted to 1/1000 in PBS-BSA (0.5%)-Triton (0.1%) before applying to samples for 1 hour. Nuclear staining was performed using Hoechst (Cayman #16756) diluted to 1/2000 in PBS-BSA (0.5%)-Triton (0.1%) for 20 minutes. Samples were washed 3 times with PBS-Triton (0.1%). Coverslips were mounted on slides using Vectashield® Antifade Mounting Medium (Vector #H-1000-10) and stored at −20°C until further acquisitions. Microscope observations were performed using a Zeiss Axio Imager 2.

### Structural and secreted GIV protein analysis

To determine the structural protein composition of GIV, GIV particles were purified as previously described^62^ and layered onto an additional sucrose gradient consisting of 30%, 40%, 50%, and 60% (w/v) sucrose at 210,000 x g to minimize interference from nonstructural proteins. The purified particles were loaded on a on a 15% SDS-PAGE for 10 minutes at 120V, followed by Coomassie blue staining to visualize the entire protein content. The whole gel was then treated by trypsin digestion followed by peptide identification by LC/MS at the Harvard Taplin Core Facility, resulting peptides were compared to the GIV proteome for identification. To determine secreted protein composition of GIV, GK cells were infected by GIV for 24h at MOI 0.1. Supernatant was harvested and supplemented with protease inhibitors, followed by 0.1μM filtration. Proteins were precipitated with addition of Trichloroacetic acid (TCA) 100% (1/5v/v) (Sigma #T6399) and washed with 0.01M HCl/ 90% Acetone before by peptide identification by LC/MS.

### Peptide synthesis

Viral insulin-like peptides were synthesized via Fmoc solid phase peptide synthesis (SPPS) utilizing a commercial automated peptide synthesizer (Symphony X, Gyros Protein Technologies). All technical details related to VILPs synthesis were previously described.

### Receptor phosphorylation and downstream phosphorylation signaling

For receptor phosphorylation and downstream signaling experiments, GK and AB9 cells were used to explore signaling properties of ligands via specific receptors. Cells were seeded into 12-well plates (100K cells per well) in 1mL of Complete Media (described above) and grown overnight. Cells were washed twice with phosphate-buffered saline (PBS) and starved in serum-free media for 4 h. After the starvation, cells were washed with serum-free media and incubated with ligand diluted in serum-free media (0, 1, 10, 100 nM) at 28°C 15 min. The reaction was terminated by washing the cells with ice-cold PBS followed by snap freezing in liquid nitrogen. Samples were stored at −80°C before further analysis.

### Western Blotting

Cellular samples analyzed by Western blot, were lysed using 50μl of RadioImmunoPrecipitation Assay (RIPA) buffer (Millipore # 20-188) supplemented with protease and phosphatase inhibitors (Selleck # B15001 & B14001). Cells on plates were incubated in the RIPA buffer on ice for 15 min, sonicated in a water bath sonicator for 1 minute and then transferred to 1.5mL Eppendorf tubes. The lysates were centrifuged (13,000 g, 5 min, 4 °C) and supernatant was transferred to new 1.5mL Eppendorf tubes. Quantification of each sample’s protein concentration were performed using the Pierce™ BCA Protein Assay Kit (ThermoFischer #23225).

Total of 6µg of protein is mixed with 6X Laemmli sample buffer (62.5 mM Tris, 2% SDS, 10% glycerol, 0.01% bromophenol blue, 0.1M DTT, pH = 6.8). Samples were loaded on a 10% SDS-PAGE gel. Proteins were transferred on PVDF Transfer Membranes, 0.45μm (ThermoFischer # 88585) using the Trans-Blot® SD Semi-Dry Transfer Cell (Bio-Rad #1703940). The membrane is then saturated in 5% BSA (GoldBio #A-420) in TBS-Tween and then probed overnight at 4C with primaries antibodies. Membrane is then washed 3 times 5 minutes with TBS-Tween. Secondary detection was performed with the Horseradish peroxidase conjugated goat anti-rabbit diluted at 1:10,000 in TBS-Tween-BSA 1% (ABclonal #AS014). Detection is made using the SuperSignal™ West Pico PLUS Chemiluminescent Substrate (ThermoFisher #34580). Acquisitions were made by exposition of autoradiography films (USA Scientific #19683810). Quantification of Western-Blot bands were made through the ImageLab software (BioRad).

### Antibodies

Antibodies used in this study are commercially available and are listed as follow: Phospho-IGF-I Receptor β (Tyr1135/1136)/Insulin Receptor β (Tyr1150/1151) (19H7) Rabbit mAb (Cell Signaling #3024); Total-IGF1R Antibody (Aviva #AVARP00004_P050), Phospho-Akt (Ser473) Antibody (Cell Signaling #9271); Akt (pan) (11E7) Rabbit mAb (Cell Signaling #4685); Phospho-p44/42 MAPK (Erk1/2) (Thr202/Tyr204) Antibody (Cell Signaling #9101); p44/42 MAPK (Erk1/2) Antibody (Cell Signaling #9102); β-Actin (D6A8) Rabbit mAb (Cell Signaling #8457); Purified anti-GFP Antibody (Biolegend # 668206)

### Replication assay and Plaque assay

Viral replication assays were performed 12 or 24 hours posts infection. Cells and supernatant were harvested followed by a snap freeze of the samples. Following the assay, viral loads were quantified by plaque assay. Briefly, samples were thawed and frozen 3 times to release all intra-cellular virions. Samples were then diluted in serial dilutions. The diluted samples were then inoculated to a confluent monolayer of GK cells placed in P12 plaques for 1 hour. Following this incubation, L-15 Media supplemented with 10% Methylcellulose was added to each well (1mL) to reduce viral dispersion during the replication. Plaques were quantified 5 days after incubation at 28°C.

### Statistics

All statistical analyses were performed using Prism6 software (GraphPad), statistical tests employed are described on each represented figure.

## Acknowledgements

We first want to thank all members of our laboratories for their valuable discussions and support. We extend our thanks to Prof. Welkin Johnson (Boston College) and his lab members for generously sharing their ultracentrifuge for our experiments. We also thank Prof. Chi-Yao Chang (Academia Sinica, Taiwan) for providing the GK cell line and GIV. Special thanks to David Pasdeloup (INRAE, France) for sharing cloning plasmids. Additionally, we acknowledge Adeline Ribeiro E. Silva for her scientific advice. We are grateful to the Novo Nordisk Research Compound Sharing Program for providing the IR Inhibitor (InsR peptide antagonist (S961)). We thank Francisco A. Valenzuela (Eli Lilly and Company) for providing dcVILP. Lastly, thanks to Bret Judson (Boston College Imaging Core) for infrastructure and support. This work is supported by the National Institutes of Health (NIH) through grant DK132674 to EA and Diabetes Research Connection grant (5116151) to AC.

