## Supplemental Figures for "Viral Insulin/IGF-like Peptides Selectively Activate Host Insulin/IGF Signaling pathways during Grouper Iridovirus Infection"

Affiliations:
[1]Boston College Biology Department, Chestnut Hill, Massachusetts, USA
[2]Department of Chemistry, Indiana University, Bloomington, Indiana, USA
[3]Department of Chemical Engineering and Bioengineering, University of New Hampshire, Durham, NH 03824, USA
[4]Department of Chemistry, University of New Hampshire, Durham, NH 03824, USA
[5]Integrated Applied Mathematics Program, University of New Hampshire, Durham, NH 03824, USA
[6]Molecular and Cellular Biotechnology Program, University of New Hampshire, Durham, NH 03824, USA

### Supplementary data

#### Figure S1

**A.**

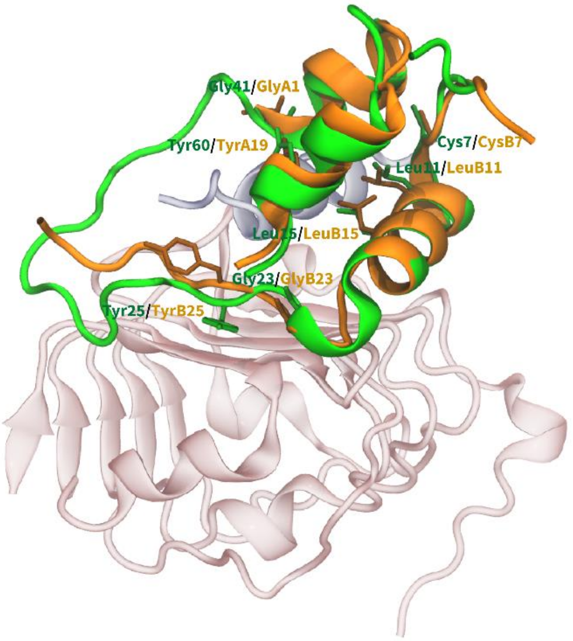


**B.**

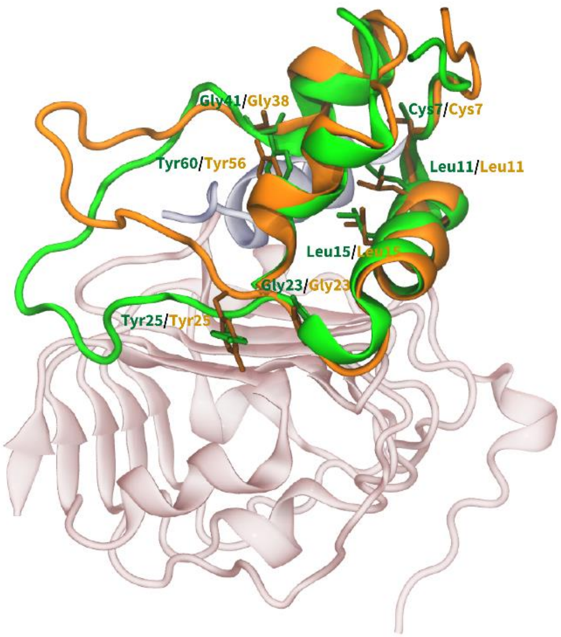


#### Figure S2

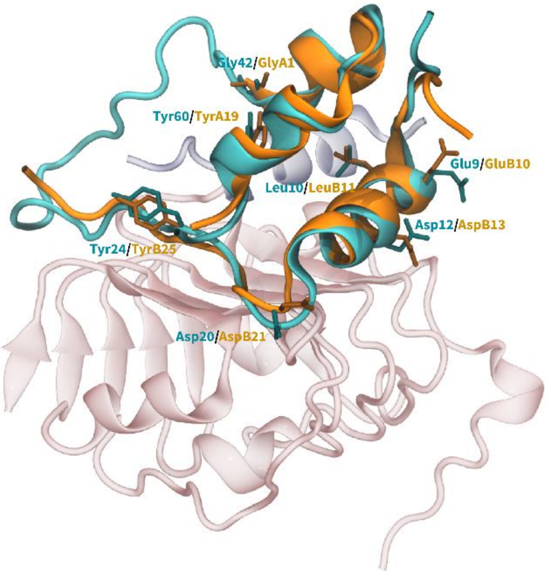

#### Figure S3

| Protein type | Signal Peptide (Sec/SPI) | Other |
| --- | --- | --- |
| Likelihood | 0.9423 | 0.0577 |

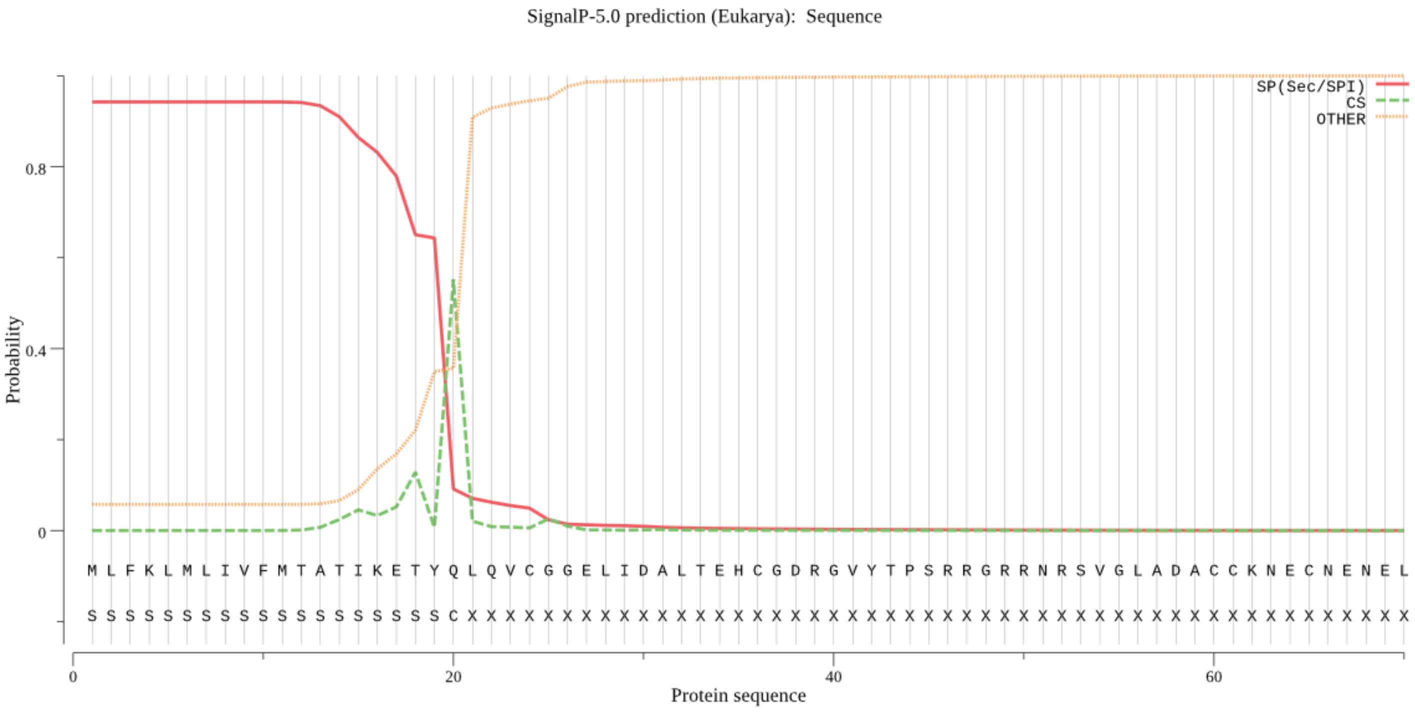


#### Figure S4

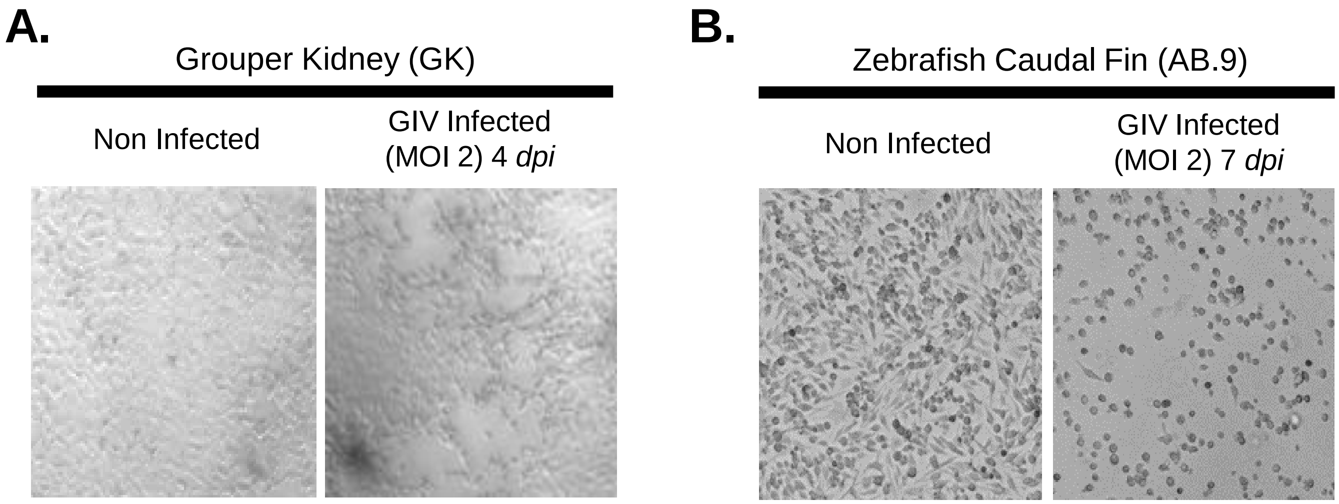
